# Mapping subarachnoid cerebrospinal fluid circulation in the human brain

**DOI:** 10.64898/2026.05.26.727780

**Authors:** Fuyixue Wang, Timothy G. Reese, Lawrence L. Wald, Bruce R. Rosen, Laura D. Lewis, Zijing Dong

## Abstract

The subarachnoid space is critical to cerebrospinal fluid (CSF) circulation and waste clearance, yet its flow organization in humans remains poorly characterized due to the lack of methods for direct, quantitative measurement of ultra-slow flow. Here we introduce slow-flow-sensitized phase-contrast imaging (SOPHI), which enables noninvasive quantification of ultra-slow flow velocities (e.g., ∼100 μm/s) and directionality at high spatiotemporal resolution, allowing whole-brain mapping of complex CSF circulation in the human subarachnoid space. We found brain-wide, spatiotemporally coherent flow patterns characterized by strong cardiac-driven oscillations. Flow dynamics were closely coupled with vascular anatomy, exhibiting higher velocities and earlier responses in subarachnoid spaces near major arteries and a spatiotemporal propagative pattern to distal spaces. We further identified localized flow pathways associated with potential CSF efflux in ventricles and key subarachnoid regions. Together, SOPHI reveals a previously undercharacterized, highly organized CSF flow system, providing a macroscopic view of human CSF circulation and a framework for investigating brain fluid transport.

## Main

The subarachnoid space plays a vital role in cerebrospinal fluid (CSF) circulation and waste clearance. The subarachnoid space forms an interconnected, brain-wide conduit that communicates with the perivascular spaces and the ventricular system, while also housing major cerebral arteries and veins. In the glymphatic model^1,2^, the subarachnoid space serves as the primary reservoir from which CSF flows into periarterial spaces and exchanges with interstitial fluid within the brain parenchyma. The subarachnoid space is also critically involved in CSF efflux pathways. Early theories propose that CSF drains from the subarachnoid space into the venous system via arachnoid granulations^3-5^. More recent studies suggest additional efflux pathways in which the subarachnoid space serves as a key compartment interfacing with these routes, including pathways along meningeal lymphatic vessels^6-8^, cranial nerves^9,10^, and bridging veins or arachnoid cuff exit^11^, further highlighting its role as a critical hub for brain-wide fluid transport. However, the CSF circulation pathways within the subarachnoid space in the human brain remain largely unexplored, owing to the limited capability of conventional imaging methods to measure slow flow in these regions. Several Magnetic Resonance Imaging (MRI) methods have been developed to probe slow CSF motion and transport, such as diffusion MRI based approaches^12-17^, spin labeling imaging and in-flow-based techniques^18-21^, and dynamic contrast-enhanced MRI^22-26^, and these techniques have advanced the physiological specificity and spatiotemporal resolution of CSF mapping. Nevertheless, these methods remain limited in their ability to provide direct, quantitative measurements of bulk flow velocity and direction, which are essential for resolving the complex flow patterns within the subarachnoid space. Phase-contrast MRI^27-29^, widely used for measuring blood flow or fast CSF flow, can directly quantify flow velocity and direction, but conventional techniques have not yet achieved sufficient sensitivity to map slow CSF flow in the subarachnoid space, and consequently, most existing studies have been limited to CSF flow measurements in the ventricles and the cerebral aqueduct.

Here, we introduce slow-flow-sensitized phase-contrast imaging (SOPHI) to elucidate brain-wide subarachnoid CSF flow patterns in the human brain. In SOPHI, we incorporate multiple acquisition, reconstruction and processing strategies to enable ultra-slow CSF flow quantification at high spatiotemporal resolution, including a slow-flow-sensitized pulse sequence, a ‘snapshot’ distortion-free readout using echo-planar time-resolved imaging (EPTI)^30,31^, a CSF-flow tailored phase-contrast processing, and a whole-brain 3D flow-vector-field visualization framework. This integrated design overcomes several limitations of conventional phase-contrast MRI that compromise sensitivity and specificity, including limited velocity encoding sensitivity for slow flow, susceptibility to physiological noise and other confounding phase sources, and image geometric distortion and its dynamic variations, while establishing a framework for visualizing and investigating brain-wide subarachnoid CSF flow.

Using SOPHI, we quantified the velocity and direction of ultra-slow CSF flow (e.g., ∼100 μm/s). We mapped brain-wide, spatiotemporal CSF flow circulation patterns in the human subarachnoid space, including both large-scale organization and localized pathways. These flows were driven by arterial pulsations and evolved rapidly over the cardiac cycle, exhibiting an oscillatory pattern with peak velocities and more rapid directional changes during systole, with overall net directional movements of substantially smaller magnitude. The cardiac-coupled flows showed a close association with vascular distribution, with greater velocities near major arteries and a spatiotemporal propagation pattern from periarterial regions to the connected peripheral subarachnoid spaces. We also observed localized pathways and directional features. For example, CSF dynamics within the lateral ventricles exhibited systolic outflow through the foramen of Monro concurrent with arterial inflow, and diastolic inflow as blood exited the brain, resulting in an overall outward net flow, consistent with the current understanding of ventricular CSF circulation^32,33^. Furthermore, we identified flow patterns that may offer new insights complementing current tracer-based and animal studies, including detailed intraventricular circulation patterns, and those in important subarachnoid regions implicated in potential CSF efflux pathways^1,3,4,11^, such as the Sylvian fissure and areas near the superior sagittal sinus (SSS). SOPHI flow measurements showed high accuracy in a slow-flow phantom with known ground-truth flow rates, as well as strong cross-subject consistency and test-retest repeatability, therefore enabling group-level characterization of brain-wide patterns. Together, these findings reveal a previously undercharacterized, highly organized CSF system within the subarachnoid space, providing a macroscopic view of bulk CSF circulation across the typical human brain and a framework for investigating brain fluid transport.

## RESULTS

### Quantification of ultra-slow CSF flow velocity and direction with SOPHI

SOPHI was designed to overcome the limited sensitivity and specificity of conventional phase-contrast imaging for measuring slow flow (**Fig. 1**). First, subarachnoid CSF flow velocities are substantially slower than those in the ventricular system, necessitating higher phase-contrast sensitivity. To achieve this, SOPHI employs a pulsed-gradient spin-echo (PGSE) sequence^34,35^ (**Fig. 1a**) in which flow-encoding gradients placed around the refocusing pulse enable a substantially longer velocity-encoding time (Δ) than conventional phase-contrast methods. This achieved significantly lower velocity-encoding (VENC) values (e.g., 1.6 mm/s vs. 100 mm/s) without increasing gradient strength or duration that would otherwise introduce unwanted diffusion weighting and CSF signal dephasing, thereby markedly improving sensitivity to ultra-slow CSF flow^36-38^ (<100 µm/s). In addition to sensitivity, specificity is also critical for accurate characterization of the measured flow signals, especially given the small phase signals of ultra-slow CSF flow. Major confounding factors include physiological noise (e.g., respiration- or motion-induced phase variations, and related image dynamic distortions), system imperfections (e.g., eddy current-induced phase variations and dynamic distortions), and contamination from blood flow. To address these at the acquisition level, SOPHI employs a single-shot spin-echo EPTI readout^30,31,39^ (**Fig. 1a**). Its single-shot acquisition avoids shot-to-shot phase inconsistencies common in multi-shot acquisitions used in conventional phase-contrast imaging, providing high physiological noise and motion robustness. At the same time, the distortion-free EPTI readout eliminates geometric distortion and its dynamic variations inherent to conventional EPI readout—effects that are particularly problematic given the narrow, confined geometry of the subarachnoid space. Furthermore, EPTI resolves multi-echo signal evolution within each readout to address contrast blurring in EPI^40^, allowing phase estimation to be restricted to spin-echo contrasts from central readout echoes, which are intrinsically less sensitive to B_0_-related phase fluctuations. Further suppression of confounding phase variations due to physiological noise or system imperfection was achieved through a dedicated CSF background phase correction framework (**Fig. 1b**), which effectively removed spatially smooth phase variations arising from respiration, motion, and eddy currents. To minimize blood-flow contributions, a long echo time (88 ms at 7T) was used to attenuate blood signals while preserving CSF signal. Residual blood-flow contributions were further suppressed through velocity encoding-induced intravoxel dephasing (effective b-value of 80 s/mm^2^), as the substantially faster blood flow results in strong intravoxel dephasing^15^. As another safeguard, voxels exhibiting large temporal phase fluctuations indicative of phase wrapping (e.g., from fast flow or noise) were excluded from phase-contrast processing, removing potential residual blood-flow signals or phase-wrapped fast flow in ventricles or large cisterns.

**Fig. 1.**
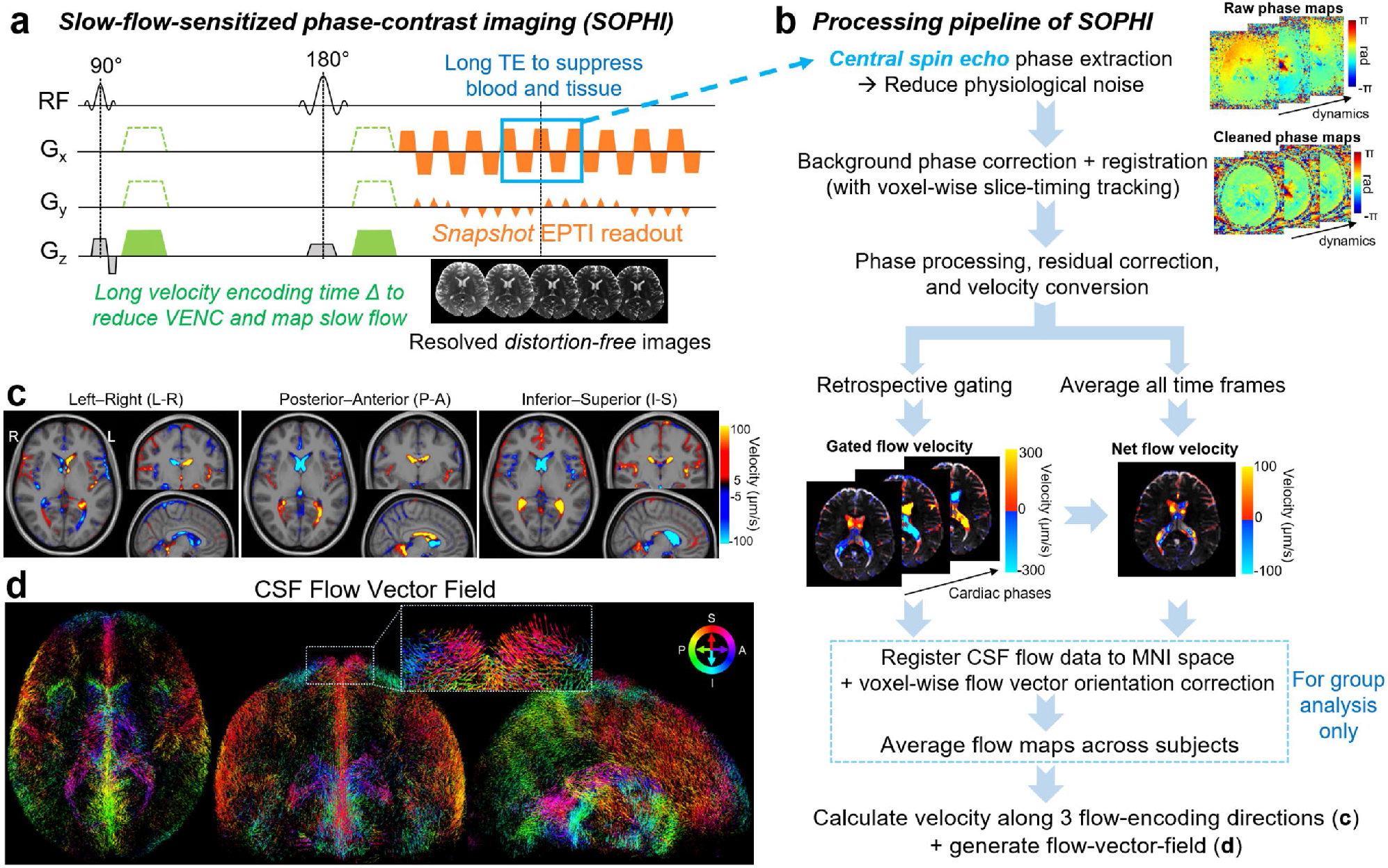
Overview of SOPHI acquisition and processing for CSF flow mapping. **a**, Illustration of SOPHI acquisition, including a pulsed-gradient spin-echo sequence with flow-encoding gradients enabling long velocity-encoding time, followed by a single-shot EPTI readout that resolves multi-echo, distortion-free images. **b**, Processing pipeline for SOPHI phase-contrast data, from raw phase images to three-dimensional CSF flow velocity and direction. **c**, Example CSF flow velocity maps of three encoding directions. Group-averaged cardiac-gated flow at cardiac phase 8 is presented. Flow-direction polarity along the three axes was defined as positive for left-to-right, posterior-to-anterior, and inferior-to-superior directions. **d**, Example CSF flow-vector-field maps (whole-volume projection of CSF voxels from different views), generated from the same data in **c**. Each voxel is represented by a flow vector, with voxel-wise vector orientation indicating flow direction, vector length proportional to 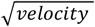, and color encoding flow direction along the I-S and P-A axes as illustrated by the color wheel at the top right (L-R is not color-coded).

We performed validation of SOPHI for slow-flow quantification using a dedicated flow phantom with controlled and known flow velocities (100–500 μm/s) and directions. Quantitative velocity measurements obtained with SOPHI were compared against the ground truth. The resulting measurements showed strong agreement with the ground truth across the tested range and small fluctuations across time frames, demonstrating high sensitivity, accuracy, and temporal stability (**Extended Data Fig. 1**).

We mapped subarachnoid CSF flow circulation patterns at the whole-brain level using 3D velocity encoding combined with 3D flow-vector-field visualization, providing both quantitative velocity and directional information. We performed group-level analysis to extract common, shared features of CSF organization across subjects. Specifically, whole-brain CSF flow data were acquired at 2-mm isotropic resolution in N = 12 healthy volunteers (31 ± 6 years; 8 females, 4 males) using SOPHI on a 7T scanner. Flow was encoded with a VENC value of 1.6 mm/s along three orthogonal directions (x, y, z). Both cardiac-gated and time-averaged flow were computed. For cardiac-gated flow, slice acquisition timing was tracked voxel by voxel even after motion registration to preserve accurate timing to match external cardiac recordings for retrospective gating. Group-level analysis was performed in MNI space following flow vector orientation correction to account for spatial transformations applied during registration from subject space to MNI space. Finally, the three directional velocity components (**Fig. 1c**) were combined to reconstruct the 3D flow-vector-field (FVF) (**Fig. 1d**) for analyzing brain-wide flow patterns.

### Brain-wide, spatiotemporal CSF flow patterns in the human subarachnoid space

We first used SOPHI to map cardiac-gated brain-wide spatiotemporal CSF flow patterns throughout the subarachnoid space. Overall, these flows evolved rapidly over the cardiac cycle (binned into 10 cardiac phases), exhibiting a spatiotemporal oscillatory pattern with more rapid directional changes (**Fig. 2a-b and Supplementary Video 1**) and elevated velocities (**Fig. 2c**) during systole, consistent with prior studies suggesting arterial pulsation as the primary driving factor of CSF flow dynamics^2,41^. Consistent with the group-averaged flow patterns (N=12) shown in **Fig. 2**, similar flow patterns were observed at the individual level and were highly consistent across subjects (individual-level flow patterns presented in **Extended Data Fig. 2**).

**Fig. 2.**
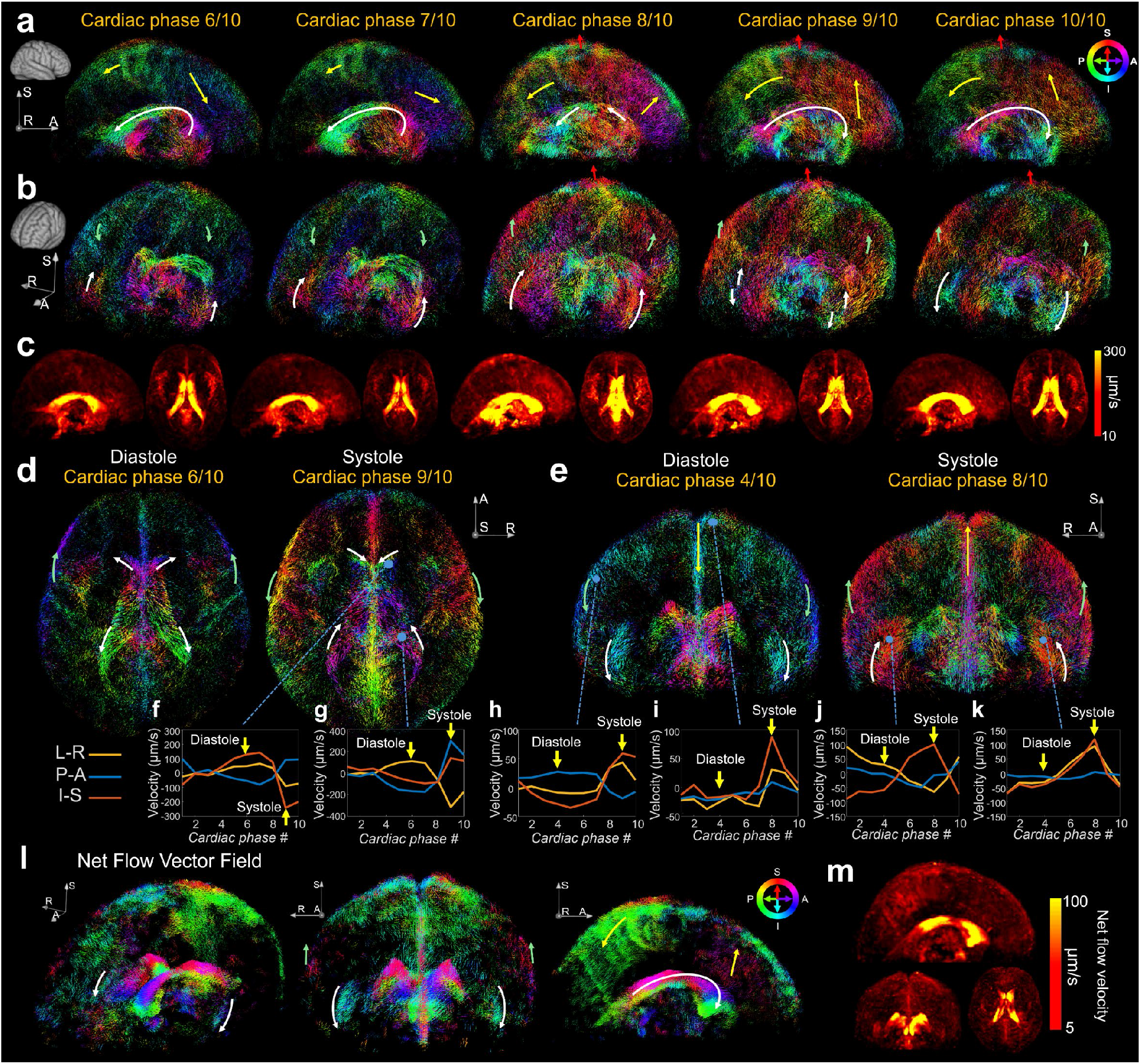
Brain-wide spatiotemporal subarachnoid CSF flow patterns. **a-b**,**d-e**, Whole-CSF-volume projections of group-averaged cardiac-gated CSF flow-vector-field maps at representative cardiac phases, shown from sagittal (**a)**, angled sagittal (**b)**, axial (**d)**, and coronal (**e**) viewing angles. Each voxel is represented by a flow vector, with voxel-wise vector orientation indicating flow direction, vector length proportional to 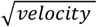, and color encoding flow direction along the I-S and P-A axes as illustrated by the color wheel at the top right (L-R is not color-coded). Flow direction can be interpreted directly from either the voxel-wise vector orientation or the color encoding, while manually drawn arrows are also included to guide visualization of the major flow patterns. White arrows: lateral ventricle and Sylvian fissure; yellow arrows: interhemispheric fissure; green arrows: outer convexity subarachnoid space; red arrow: subarachnoid space near the SSS. **c**, Maximum intensity projections of group-averaged cardiac-gated CSF velocity maps at cardiac phases 6-10. **f-k**, Time series of cardiac-gated CSF flow velocities (along three directions) across all ten cardiac phases from voxels in representative regions marked by blue dots in **d-e. l**, Whole-CSF-volume projections of group-averaged net CSF flow-vector-field maps from different views, illustrating distinct flow patterns summarized by colored arrows. **m**, Maximum intensity projections of group-averaged net CSF flow velocity maps.

Representative large-scale flow pathways obtained by SOPHI are illustrated in **Fig. 2a-b,d-e** (marked by arrows) with corresponding velocity time series across cardiac cycle in **Fig. 2f-k**. In general, cardiac-gated phases 3–7 typically corresponded to diastole and phases 8–10 to systole (**Fig. 2a-b**); however, the timing and phases of systole/diastole vary across different regions. Flow direction can be interpreted directly from either the voxel-wise vector orientation or the color encoding as indicated by the color wheel, while manually drawn arrows are also presented to guide visualization of the major flow patterns. Specifically, for example, within the lateral ventricles (**Fig. 2a,d**; white arrows), CSF was observed to flow inward from the third ventricle during diastole (e.g., cardiac phase 6-7) and reverse during systole (e.g., cardiac phase 9-10), consistent with the well-established observations in prior studies^32,33^ reporting diastolic CSF inflow due to cerebral blood outflow and systolic CSF outflow as blood enters the brain. The direction reversal is also evident in the velocity time series (**Fig. 2f,g**), where the velocities changed sign at the diastole-systole transition. Similar to ventricular flow, subarachnoid flows exhibited oscillatory dynamics across the cardiac cycle. For example, CSF within the interhemispheric fissure (**Fig. 2a,e**; yellow arrows) and the outer convexity subarachnoid space (**Fig. 2b,d-e**; green arrows) showed flow velocity and direction changes between diastole and systole. Example time series from the anterior outer convexity subarachnoid space (**Fig. 2h**) demonstrated upward peak velocities at cardiac phase 9. In the lateral Sylvian fissure (**Fig. 2b,e**; white arrows), coherent upward flow was observed during systolic cardiac phases (e.g., phase 7-8), and reversed to downward flow afterwards. Notably, compared with other previously mentioned regions, the Sylvian fissure exhibited an earlier systolic peak. Velocity time series (**Fig. 2j,k**) confirmed this pattern, characterized by their strong z-directional flow being upward during phases 6–8 and downward during phase 10,1–4, indicating regional differences in the temporal responses of subarachnoid CSF flow to arterial pulsation. As a localized feature, systolic upward flow was observed near the superior portion of the SSS (**Fig. 2b,d-e**; red arrows) and confirmed by the time series (**Fig. 2i**). Overall, the flow patterns exhibited coherent directions aligned with the underlying anatomical curvature. Left–right symmetry was observed in both the flow-vector-fields and velocity time series across regions. For example, the left and right Sylvian fissures exhibited opposite left-right velocity directions consistent with anatomical expectations; similar symmetry was also observed in other regions, including the ventricles.

In addition to cardiac-gated flow that reflects transient flow dynamics, we also calculated net flow velocity (i.e., time-averaged mean velocity) to reflect cumulative CSF transport within the subarachnoid space (**Fig. 2l**). Net flow velocities derived from both time averaging without cardiac-gating, and from time averaging across the cardiac-gated time series were computed, which yielded similar results (**Extended Data Fig. 3**). Compared with cardiac-gated flow, net flow velocities were substantially smaller (**Fig. 2m**; typically below 100 μm/s), suggesting that CSF flow was dominated by cardiac-driven oscillatory dynamics with a small net transport component. Given the low magnitude of net flow velocities, we focused on large-scale, coherent flow patterns that were consistent across subjects (corresponding individual subject maps shown in **Extended Data Fig. 4**). Overall, the cumulative CSF flow data were noisier than the cardiac-gated flow but still revealed coherent brain-wide flow patterns with local heterogeneity across the subarachnoid space. Specifically, net flow in the lateral ventricles was consistent with prior reports^32,33^ showing posterior-to-anterior outflow toward the third ventricle (**Fig. 2l**; white arrow). CSF within the lateral Sylvian fissures exhibited predominantly inferiorly directed net flow (**Fig. 2l**; lateral white arrows). Within the interhemispheric fissure, the anterior segment showed superiorly directed flow, whereas the posterior segment exhibited posterior-inferior flow (**Fig. 2l**; yellow arrows). In the anterior regions of convexity subarachnoid spaces, superiorly directed flow was observed (**Fig. 2l**; green arrows).

### Association between subarachnoid CSF flow propagation and vascular anatomy

Given the strong cardiac-coupled CSF flow responses observed, we next examined the spatial distribution and temporal characteristics of flow dynamics, as well as their association with vascular anatomy (**Fig. 3**). We observed high velocities in the ventricles and in subarachnoid regions adjacent to major arteries (**Fig. 3a,b**), supported by strong spatial correspondence between the CSF velocity maps and the vessel probability atlas^42^ (**Fig. 3c**). These regions include the Sylvian fissure proximal to the middle cerebral artery (MCA), the quadrigeminal cistern containing the posterior cerebral artery (PCA), and the anterior interhemispheric fissure, potentially reflecting the influence of pericallosal artery pulsation. In addition to these periarterial regions, subarachnoid flow near the SSS also exhibited high velocities. Quantitatively, periarterial regions around MCA and PCA exhibited high velocity ranges (defined as the difference between maximum and minimum velocities across the cardiac cycle for each encoding direction and combined across all three directions), reaching approximately 500 μm/s (**Fig. 3a**), along with high peak absolute velocities of approximately 300 μm/s (**Fig. 3b**). Regions around the pericallosal artery and SSS also showed high velocity ranges and peak absolute velocities, albeit lower than those observed around the MCA and PCA.

**Fig. 3.**
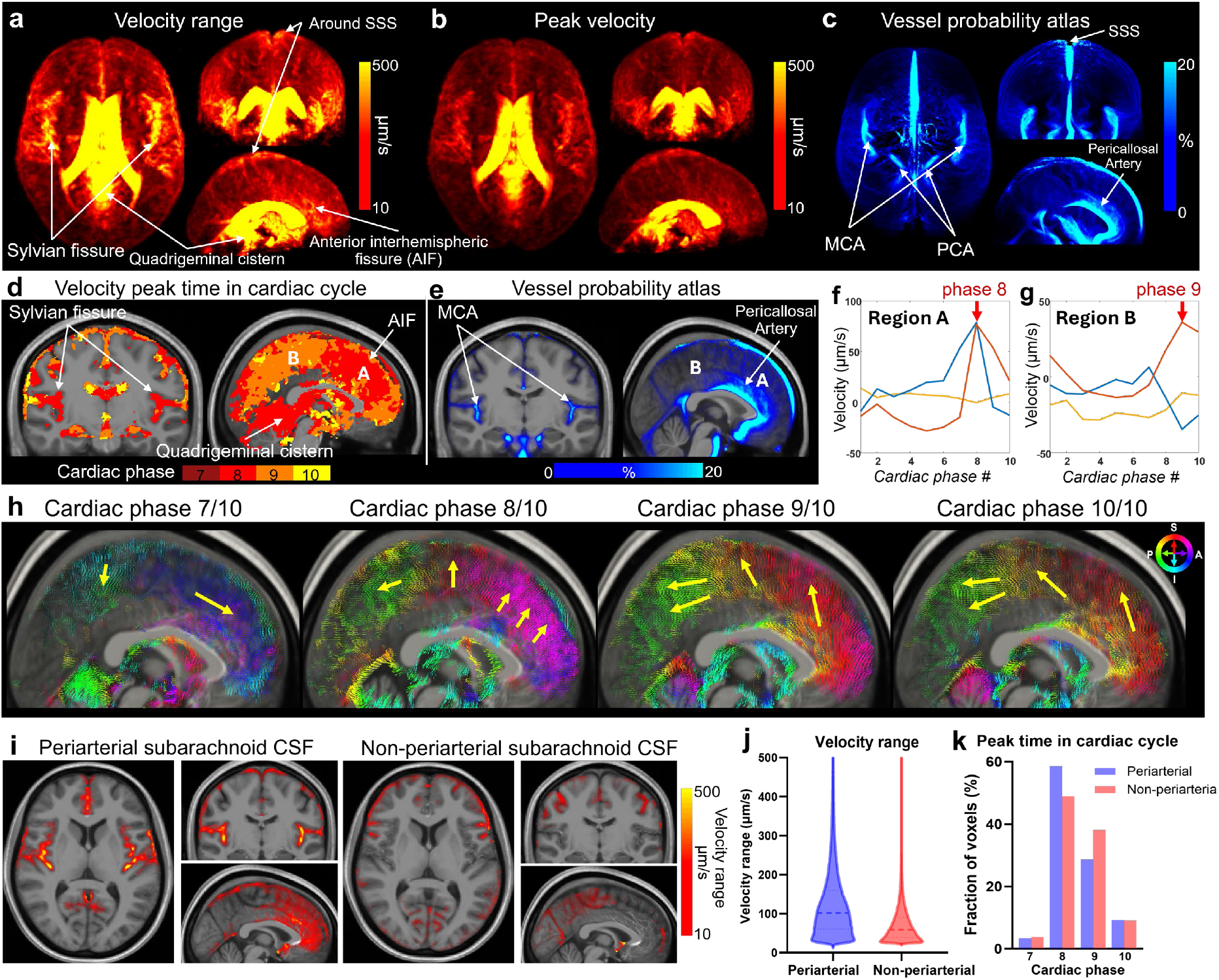
Spatiotemporal characterization of cardiac-gated subarachnoid CSF flow and their association with vascular distribution. Maximum intensity projections of group-averaged velocity range (**a**) and peak velocity (**b**) of the cardiac-gated CSF flow. **c**, Maximum intensity projections of vessel probability atlas obtained from Magnetic Resonance Angiography, primarily representing arterial distribution with partial inclusion of venous sinus structures. **d**, Maps of velocity peak time within cardiac cycle, defined as the cardiac phase at which each voxel reaches its peak velocity. **e**, Vessel probability atlas. **f-g**, Time series of cardiac-gated CSF flow velocities (along three directions) from example voxels in region A and B, with red arrows indicating peak cardiac phase. **h**, Flow-vector-field maps illustrating flow direction changes and their spatiotemporal propagation across cardiac phases. A thin midsagittal slab within the interhemispheric fissure is shown instead of the whole-CSF-volume projection. Voxel-wise vector orientation indicates flow direction, vector length is proportional to 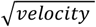, and color encodes flow direction along the I-S and P-A axes as illustrated by the color wheel at the top right. Yellow arrows summarize the flow patterns. **i**, Velocity range maps segmented into periarterial and non-periarterial subarachnoid CSF space. **j**, Violin plots of velocity ranges within periarterial and non-periarterial subarachnoid CSF spaces, with medians indicated by thick dashed lines and quartiles indicated by thin dashed lines. **k**, Histogram of velocity peak time across the cardiac cycle for periarterial and non-periarterial subarachnoid CSF flow.

In terms of temporal characteristics, the velocity peak-time map in **Fig. 3d** suggested that peak velocities occurred earlier in subarachnoid regions located near major arteries (red denotes earlier cardiac phases)—including the Sylvian fissure, quadrigeminal cistern, and anterior interhemispheric fissure—indicating faster temporal responses to arterial pulsation. Using the interhemispheric fissure as an example, the anterior segment (region A), adjacent to the pericallosal artery, reached peak velocity at phase 8, while the posterior segment (region B) peaked at phase 9 (∼100 ms delay compared to phase 8), as shown in both the velocity peak-time maps (**Fig. 3d**) and the velocity time series (**Fig. 3f,g**). This propagation of flow can also be observed in the flow-vector-field maps (**Fig. 3h**): from cardiac phase 7 (diastole) to cardiac phase 8 (systole), flow direction of the anterior segment changed from downward-anterior to upward-anterior and the velocity intensity reached peak, likely driven by the pulsation of pericallosal artery; at phase 9, flow velocities in the anterior segment started to decrease and changed to an upward-posterior direction, while the posterior segment exhibited accelerated backward flow and reached its velocity peak; all velocities subsequently decreased after phase 10. This propagation pattern was further resolved with finer temporal sampling using cardiac gating with 20 phases (**Extended Data Fig. 5**). These results highlight the interconnected and propagative nature of the subarachnoid CSF flow driven by arterial pulsations, forming large-scale structured flow patterns.

By segmenting the periarterial (near large major arteries) and non-periarterial subarachnoid spaces (**Fig. 3i**), we found that periarterial CSF flow exhibited a higher velocity range (**Fig. 3j**; median value 101.65 μm/s vs. 59.10 μm/s, p < 0.0001) and a greater proportion of voxels with earlier peak timing (**Fig. 3k**) compared with the non-periarterial space. These findings underscore the correspondence between arterial distribution and subarachnoid CSF flow spatiotemporal dynamics.

### Localized CSF flow patterns and pathways in regions associated with CSF efflux

We next examined localized CSF flow patterns (**Fig. 4**) in key regions potentially associated with CSF efflux, including the better-understood regions such as the ventricles for validation and in less-explored subarachnoid regions. In the lateral ventricles (**Fig. 4a**), CSF flowed outward through the foramen of Monro during systole as blood entered the brain and flowed inward during diastole (indicated by the large white arrows, which is based on the voxel-wise vector orientations), consistent with the established understanding^32,33^. Notably, additional intraventricular patterns beyond prior reports were observed: near end-systole, the outflow decreased and partial backflow emerged, forming a localized circulating pattern in the inferior portion and posterior horn of the lateral ventricles (orange arrows). This behavior may reflect a reduction in the pressure gradient driving ventricular outflow as cerebral blood inflow subsides toward end-systole, causing part of the outflow to be pushed back and generate intraventricular circulation. Consistent with this observation, the net-flow pattern of the lateral ventricle resembles the end-systolic pattern, showing overall outward flow transport accompanied by intraventricular circulation. In the Sylvian fissure (**Fig. 4b**), CSF flowed upward during systole and downward during diastole (white arrows), with an overall downward net flow, likely entering the basal cisterns connected to the Sylvian fissure and subsequently exiting the brain. Another distinct pattern was observed near the superior portion of SSS (**Fig. 4c**), where upward flow from the adjacent subarachnoid space on both the left and right sides converged toward the midline SSS during systole (**Fig. 4c**, top row), with substantially lower velocities during diastole. Around the relatively posterior portion of the SSS (**Fig. 4c**, bottom row), subarachnoid CSF flowed posteriorly toward the posterior SSS. The resulting net flow resembled the systolic pattern but with much lower amplitude. These observations are consistent with prior studies suggesting potential CSF drainage via arachnoid granulations near the SSS^3-5,43^. The regional flow patterns reported here were also at the group-averaged level and were consistent across the scanned healthy subjects. Specifically, all subjects exhibited the lateral ventricular and Sylvian fissure flow patterns, along with clear systolic upward flow near the SSS (example individual flow maps in **Extended Data Fig. 2** and **4**).

**Fig. 4.**
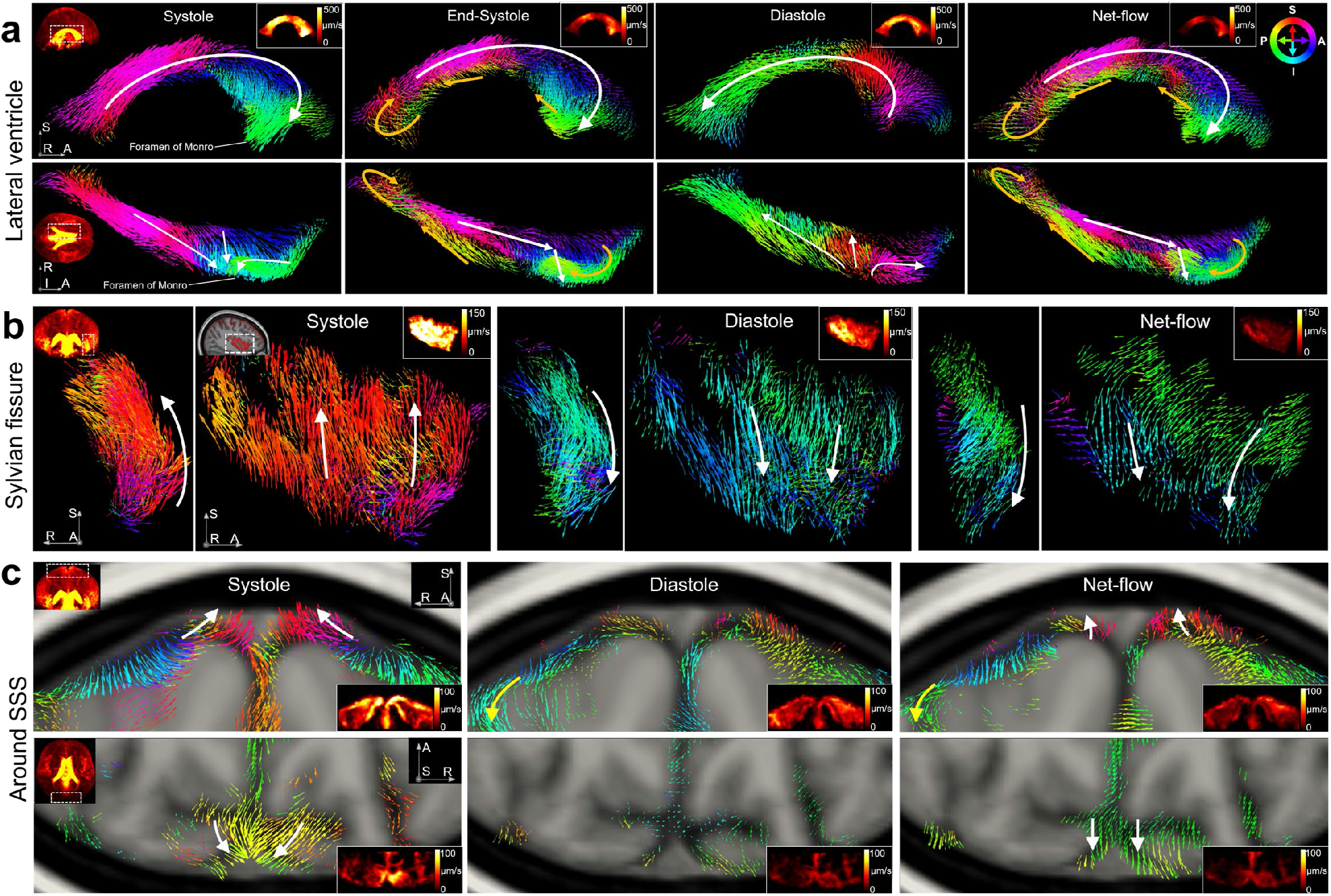
Localized CSF flow patterns and pathways. Group-averaged systolic, diastolic, and net CSF flow-vector-field maps, shown as region-specific projections from multiple views. Voxel-wise vector orientation indicates flow direction, vector length is proportional to 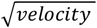, and color encodes flow direction along the I-S and P-A axes as illustrated by the color wheel at the top right. Insets to the left of the first column indicate the locations of the zoomed-in regions (white dashed boxes). Insets at the top or bottom right of each panel show maximum-intensity projections of flow velocity within the corresponding regions. **a**, Lateral ventricle. Top and bottom rows show the sagittal and axial (inferior-to-superior) views, respectively. White arrows indicate the overall flow direction, and orange arrows indicate backflow and intraventricular circulation. **b**, Sylvian fissure. Within each column, the left and right panels show the coronal and sagittal views. White arrows indicate the overall flow direction. **c**, Regions near the SSS. Top row shows coronal slabs of the superior portion of SSS, and bottom row shows axial slabs near the posterior portion of SSS. White arrows show the overall flow direction near the SSS.

### Validation and repeatability characterization of flow measurements

In addition to the flow-phantom validation mentioned above, we assessed the in-vivo reliability of subarachnoid CSF flow measurements acquired using SOPHI in 12 healthy volunteers. Scan–rescan acquisitions were performed for each subject, and quantitative metrics were evaluated across 70 cortical-based subarachnoid ROIs (**Fig. 5**).

**Fig. 5.**
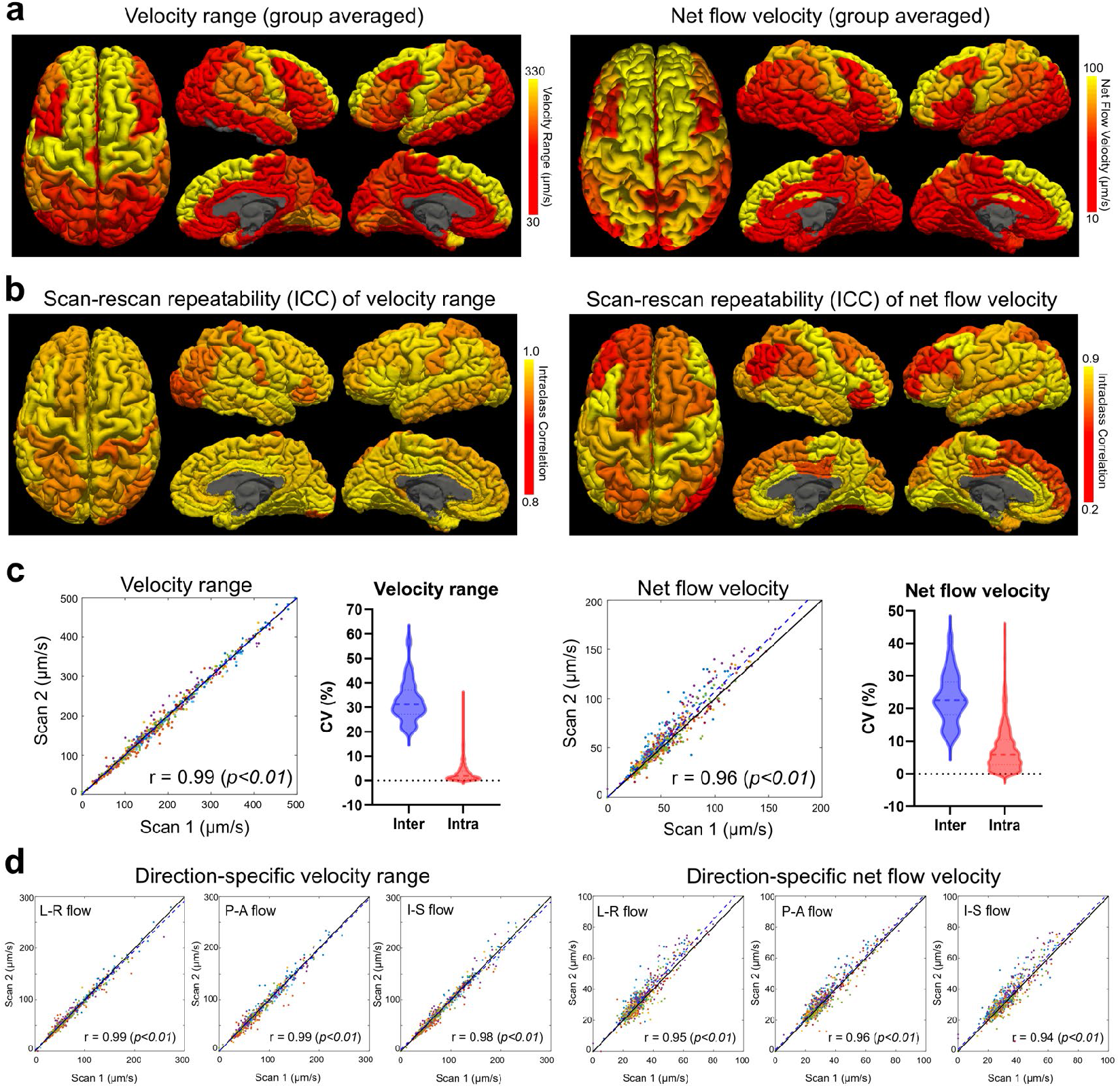
ROI-based CSF flow characterization and scan-rescan repeatability analysis. **a**, Group-averaged, ROI-based velocity range over the cardiac cycle (left panel) and net flow velocity (right panel) overlaid on brain surfaces. **b**, Intraclass correlation coefficients (ICC) for velocity range (left panel) and net flow velocity (right panel), assessing scan-rescan repeatability. **c**, For velocity range (left panel) and net flow velocity (right panel), scatter plots show scan-rescan repeatability across all ROIs and participants, with different colors denoting individual subjects. The fitted regression line (dashed blue) is shown alongside the identity line (black). Pearson correlation coefficient (PCC, r) is shown at the bottom-right of each plot. Violin plots of the coefficient of variation (CV) are also shown, indicating inter-subject and intra-subject variability, with median indicated by the thick dashed line and quartiles indicated by thin dashed lines. **d**, Scatter plots of scan-rescan repeatability of direction-specific velocity range and net flow velocity for the three flow-encoding directions.

We first evaluated the scan-rescan repeatability of flow velocity magnitude using the velocity range over a cardiac cycle and the net flow velocity. Regions surrounding the superior frontal gyrus and precentral gyrus exhibited relatively higher velocities in both velocity range and net flow velocity (**Fig. 5a**). For velocity range, all subarachnoid ROIs demonstrated excellent scan– rescan repeatability, with intraclass correlation coefficient (ICC) values higher than 0.9 (**Fig. 5b**, left panel). For net flow velocity, most ROIs showed moderate to high repeatability, with ICC values greater than 0.6 (**Fig. 5b**, right panel). In addition, scan-rescan measurements exhibited strong correlations with high Pearson correlation coefficients (PCC) values for both velocity range (PCC = 0.99, p < 0.01, **Fig. 5c** left panel) and net flow velocity (PCC = 0.96, p < 0.01, **Fig. 5c** right panel), further supporting measurement repeatability. In addition to the direction-combined velocity, we also assessed direction-specific repeatability for velocities along each flow-encoding direction (**Fig. 5d**). Velocity ranges across all three flow-encoding directions showed high repeatability with PCC values exceeding 0.98, and net flow velocities across all directions similarly exhibited strong scan-rescan correlations (PCC > 0.94). These analyses indicate that cardiac-pulsation-driven CSF flow magnitudes exhibited high values and excellent repeatability, while net flow velocities, despite their lower magnitudes, also demonstrated good repeatability. In addition, low intra-subject variability but high inter-subject variability was observed for both metrics, with pulsation-driven velocity range showing greater inter-subject variations than net flow velocity (**Fig. 5c**; mean coefficient of variation: 32.4% vs. 23.2%).

## Discussion

The present study developed SOPHI—an MRI acquisition technology with integrated processing and visualization frameworks— to characterize CSF flow in the human subarachnoid space, a highly complex and anatomically heterogeneous compartment critical to brain-wide CSF circulation and waste clearance. The results obtained by SOPHI revealed that subarachnoid CSF flow exhibits coherent large-scale pathways accompanied by localized patterns, which are spatially interconnected and temporally propagating, and strongly shaped by the complex geometry of the subarachnoid space and the underlying cerebral vasculature. The results suggest that subarachnoid CSF flow system is highly structured with organized flow patterns dominated by cardiac-coupled oscillatory motion accompanied by modest directional net movements.

The observed brain-wide flow patterns—particularly their oscillatory behavior across the cardiac cycle and close association with arterial distribution—highlight the important role of arterial pulsation in driving transient CSF dynamics. Our results further revealed a propagative pattern of the flow, in which periarterial regions respond earlier with greater magnitude, followed by delayed responses in interconnected subarachnoid spaces. This observation may support a dynamically coupled CSF system, in which arterial pulsation induces local CSF displacements that in turn generates a pressure gradient and propagates into anatomically connected spaces, therefore producing globally coordinated flow dynamics. The observed high velocities and more rapid temporal responses in the periarterial subarachnoid space may support previous reports of faster enrichment or transport of CSF tracers in these regions^25^. The flow-related pressure gradient and larger-scale propagation can further facilitate CSF mixing or tracer transport over longer distances^44^. The close correspondence between the spatiotemporal characteristics of subarachnoid CSF flow and arterial distribution indicates the importance of vascular anatomy and function for CSF circulation.

Beyond global organization, SOPHI revealed localized flow pathways that not only provide confirmation of more established features of CSF circulation, but also offer new insights into patterns potentially linked to CSF efflux pathways under active investigation, consistent with and complementing prior tracer-based and animal studies^4^. For example, ventricular flow patterns recapitulate known inflow–outflow dynamics while additionally suggesting intraventricular circulation, indicating more complex internal mixing and redistribution processes. The systolic convergent upward flow toward the SSS, with consistent spatial distribution with recent contrast-enhanced studies suggesting tracer efflux to the parasagittal dura in humans^43^, may be related to CSF absorption through arachnoid granulations^3-5^ and/or the arachnoid cuff exit points^11^. Similarly, the net downward flow in the Sylvian fissure may suggest transport toward basal cisterns and downstream drainage routes. While these findings delineate the outer spaces/routes in the subarachnoid space and its macroscopic circulation, we observed downward systolic and net flow along many of the cortical sulcal regions (**Fig. 4c**; yellow arrows), which might be associated with the coupling between the subarachnoid CSF flow and perivascular spaces, motivating further investigation. An important observation was that net CSF flow velocities were substantially smaller than the cardiac-driven oscillatory flow, indicating that bulk CSF transport may arise from subtle imbalances in large bidirectional pulsatile flow rather than sustained unidirectional flow—a similar effect has been previously observed in optical imaging studies in rodents^41^. These findings have important implications for interpreting CSF circulation: its transport mechanisms are likely sensitive to physiological and pathological changes that alter vascular pulsatility, compliance, and/or the boundary conditions of the subarachnoid space.

The validity of the flow measurements was assessed through multiple complementary approaches. First, accuracy was established using a slow-flow phantom with known ground truth, demonstrating reliable quantification under controlled conditions. For in vivo measurements, which are subject to additional confounds such as physiological noise and residual phase artifacts, we evaluated both scan–rescan repeatability and cross-subject consistency. Scan–rescan analyses demonstrated good reproducibility of pulsation-driven velocity range and flow directionality, as well as good reproducibility of net flow velocity. For cross-subject consistency, major flow pathways identified were consistently reproduced across subjects, arguing against artifactual origins such as motion or physiological noise, which would be expected to vary idiosyncratically across individuals. Beyond these shared patterns, inter-subject heterogeneity was observed at the same time, especially in the higher intensity cardiac-driven flow, which is in line with expectations given the large individual variabilities in anatomy, CSF volume, subarachnoid geometry, and vascular physiology. Finally, the observed spatiotemporal coherence of the flow patterns provides additional support for measurement validity. The flow pathways exhibited structured spatial organization aligned with the underlying anatomy, including the observation of left–right symmetry. They also demonstrated coherent temporal responses after data resorting in cardiac gating. These spatially and temporally coherent, and physiologically coupled patterns—together with their propagation across connected regions—are unlikely to arise from artifacts. Moreover, the net flow is spatially concordant with the oscillatory flow features and shows similar localized patterns, further supporting their physiological origin.

The high sensitivity and acquisition efficiency of SOPHI enables rapid whole-brain mapping of subarachnoid CSF flow (e.g., 3 s per volume, three-directional velocity, each with 80 dynamics, total scan time 12 minutes). This fast speed makes SOPHI well suited for clinical studies, particularly for conditions associated with altered CSF circulation, such as impaired clearance in Alzheimer’s disease^45-47^ and disrupted flow pathways in normal pressure hydrocephalus^48-50^. Although the present study was performed at 7T, translation to lower field strengths (e.g., 3T) is readily feasible. To preserve the sensitivity and specificity at lower field strengths, SOPHI can be implemented with similar velocity-encoding parameters but increased echo times to enhance T_2_ weighting and attenuate blood signals, at the cost of modestly increased TR and scan duration. Implementation at lower field strengths also offers complementary advantages, including improved B^1^ homogeneity for investigating brainstem and cervical regions and reduced physiological noise. This would enhance accessibility for clinical and translational studies and position SOPHI as a promising tool for investigating disease-associated alterations in CSF circulation, advancing mechanistic understanding and evaluation of therapeutic interventions.

Finally, we outline several directions for future work that highlight the potential impact of this technique. First, while the current work primarily depicted CSF flow dynamics in the subarachnoid space and ventricles, future extensions can map flow pathways across the perivascular space and the broader ventricular system, enabling an integrated, multi-scale view of brain-wide CSF circulation and inter-compartmental connectivity. In particular, incorporating a multi–velocity-encoding (multi-VENC) strategy^51,52^ would provide sensitivity to both ultra-slow (subarachnoid and perivascular space) and faster CSF flow components (e.g., fourth ventricles). Combined with further gains in spatial resolution achievable with SOPHI and distortion-free EPTI readouts, this may allow direct visualization of fine-scale perivascular flow pathways in the living human brain. Beyond arterial pulsation, SOPHI provides a useful tool to investigate other CSF driving factors (e.g., respiration^18,53,54^, brain activity^19,55,56^) and their associated spatiotemporal flow patterns, including neural activity-related ultra-slow flow changes on the order of ∼20 μm/s^37,57,58^. Moreover, all observations in the present study were obtained in the supine position. Because body posture is known to influence CSF dynamics^59,60^, future studies will investigate how flow organization varies with body positions. Finally, applying SOPHI to aging populations and neurological disorders may provide important insights into how alteration in both brain-wide and region-specific CSF flow circulation contributes to impaired waste clearance, glymphatic dysfunction, and disease progression.

To summarize, our findings establish a novel framework for noninvasively mapping brain-wide and regional CSF flow spatiotemporal organization, propagation, and directional transport in the human brain, together with their relationships to anatomy and physiology. By enabling quantitative characterization of CSF flow magnitude and directionality, SOPHI opens new opportunities for investigating the physiological mechanisms, functional roles, and pathological alterations of the CSF circulation system in health and disease.

## METHODS

### SOPHI acquisition and sequence design

SOPHI employs a PGSE sequence^34,35^ with EPTI readout^30,31,39^ to acquire phase-contrast data with high sensitivity and specificity to slow CSF flow (**Fig. 1a**). In PGSE, the flow-encoding gradients before the readout encode spin displacement along a specified spatial direction, generating velocity-dependent phase shifts in the sequential readout signals. A key advantage of PGSE-based flow encoding over conventional gradient-echo phase-contrast methods is its ability to achieve substantially longer velocity encoding time (Δ) without excessive signal loss (e.g., due to strong T2* decay in conventional gradient echoes) or confounding phase accumulation (e.g., B^0^ phase accumulated across gradient echoes), enabling lower VENC values and thereby improving sensitivity to slow flow. Increasing Δ also allows reduction of VENC without requiring larger gradient amplitudes (G) or durations (δ), which would otherwise increase CSF signal dephasing and reduce signal-to-noise ratio (SNR)^35,38^ due to diffusion weighting (i.e., b-value) that scales only linearly with Δ but quadratically with G and δ. Note that the VENC value defines the maximum velocity that can be measured by phase-contrast MRI without inducing phase wrapping. In this study, a VENC value of 1.6 mm/s was used for all experiments, corresponding to a b-value of 80 s/mm^2^, which preserved sufficient SNR for CSF signals.

To improve measurement specificity, SOPHI was designed to minimize confounding factors from blood signal and physiological noise. A relatively long echo time (TE = 88 ms) at 7T was employed to enhance T^2^ weighting and suppress blood signal, which exhibits much shorter T_2_ relaxation times than CSF, and residual blood signals are further minimized by the strong dephasing effect of the velocity-encoding gradients (effective b-value of 80 s/mm^2^) on fast-flowing spins^15^. The strong T^2^ weighting also suppresses parenchymal signal, thereby reducing partial volume effects on phase contrast at the tissue–CSF interface. For physiological noise, major sources include respiration- and motion-related phase variations, especially motion occurred during velocity encoding. Conventional gradient-echo phase contrast imaging typically uses multi-shot acquisition, making it inherently more vulnerable to shot-to-shot phase-induced physiological noise. Although single-shot EPI can mitigate this vulnerability, it suffers from B_0_-induced image geometric distortion, and B_0_ fluctuations will further lead to temporal variations in image geometry, particularly problematic when mapping small subarachnoid space. To address these limitations, we employed EPTI as the imaging readout in a single-shot regime^30^ (**Fig. 1a**), which eliminates geometric distortion and its temporal variations while avoiding shot-to-shot phase inconsistency and reducing vulnerability to motion and physiological noise, thereby improving temporal stability and overall sensitivity. Moreover, EPTI recovers the full k-TE space and resolves multi-echo images within the readout, rather than collapsing all signals across the readout to form a single image with image/contrast blurring as in EPI. In this study, 95 echoes were reconstructed within EPTI readout, and only the central 28 echoes (∼29%) around the spin-echo TE, which are inherently less sensitive to B_0_ -related fluctuations, were used for phase-contrast reconstruction, providing further improved robustness to respiration-related physiological noise.

### Experimental design and data acquisition

All volunteers provided written informed consent prior to participation under an Institutional Review Board (IRB)–approved protocol, in accordance with the policies of our institution’s Human Subjects Research Committee. Participants were healthy adults (N = 12, 31 ± 6 years; 8 females, 4 males). Experiments were performed on a whole-body 7T MRI system (MAGNETOM Terra, Siemens Healthineers, Erlangen, Germany) using a 32-channel head array coil (Nova Medical, Wilmington, MA, USA) with participants being in supine position. Cardiac physiological signals were recorded concurrently during all scan sessions using a fingertip piezoelectric device.

For both the slow-flow phantom experiments and in vivo studies, the same SOPHI–EPTI protocol was employed: whole-brain phase-contrast data were acquired at 2-mm isotropic resolution with a VENC value of 1.6 mm/s. Imaging parameters included: FOV = 216 × 216 × 92 mm^3^, TR/TE = 3000/88 ms, multiband factor = 2, echo spacing = 0.6 ms, 95 EPTI echoes with the spin echo at the center (TE = 88 ms), no partial Fourier, and an effective b-value of 80 s/mm^2^. For in vivo experiments, each run consisted of three orthogonal velocity-encoding directions (x, y, z) acquired sequentially, with 80 dynamics per direction (six non-velocity-encoded calibration volumes and seventy-four velocity-encoded volumes for phase-contrast calculation), resulting in a total acquisition time of approximately 12 minutes for all three directions (3 s per dynamic volume x 80 dynamics per direction x 3 directions). The three velocity-encoding directions were acquired in world coordinates, independent of slice orientation. A fast, low-resolution EPTI calibration pre-scan (∼50 s) was acquired before each run to estimate coil sensitivity and B^0^ field maps for subspace reconstruction. For test–retest repeatability assessment, two repeated measurements were obtained within the same scan session for each subject, each consisting of two runs of the three-directional acquisition (in total of ∼24 minutes).

Except for repeatability analyses, flow maps presented in this study were generated using all available runs (four runs per subject) to maximize SNR. For the slow-flow phantom experiments, 30 dynamics were acquired with z-direction velocity encoding, including six non-velocity-encoded calibration volumes and 24 velocity-encoded volumes. Anatomical reference images for brain segmentation were acquired using a standard T^1^-weighted MPRAGE sequence at 0.75-mm isotropic resolution. The vessel probability atlas was derived from an existing open Magnetic Resonance Angiography (MRA) dataset of healthy subjects^42^, which predominantly represents arterial distribution, with some inclusion of venous sinuses.

### SOPHI data reconstruction and processing

Standard EPTI reconstruction using a subspace approach^30,61,62^ was performed in MATLAB (MathWorks, Natick, MA, USA) and BART^63,64^ to reconstruct images from raw k-space data. Coil sensitivity maps were estimated using ESPIRiT^65^ to ensure accurate phase reconstruction. After image reconstruction, phase-contrast data processing was employed in SOPHI to generate three-dimensional velocity and flow-vector-field maps from the phase images (**Fig. 1b**). For each time frame, central echo images were first averaged to obtain a single phase image, followed by background phase correction using a third-order spatial polynomial fitting on each slice, excluding CSF voxels, to estimate and remove spatially smooth background phase variations arising from respiration, motion, or eddy currents. Flow-induced phase, which is spatially higher frequency and more localized, was minimally affected by this procedure. Then, rigid motion correction was performed using AFNI^66,67^ with motion parameters estimated from all-echo-averaged magnitude images and subsequently applied to the phase volumes. Nearest-neighbor interpolation was used here to preserve accurate voxel timing—acquisition timing for each voxel was tracked by accounting for their original slice-timing even after motion correction re-interpolation and was later used for retrospective cardiac gating. Following registration, phase differences between each velocity-encoded volume and non-velocity-encoded calibration volume were calculated, followed by applying a second third-order spatial polynomial fitting to further remove residual background phase. Phase values were subsequently converted to velocity according to velocity = (phase/π) × VENC. The non-velocity-encoded calibration volumes were acquired at the beginning of each run and averaged across multiple time frames to reduce physiological variability. CSF masking was applied to the velocity maps by combining an intensity-based CSF mask with a CSF segmentation mask obtained using the FAST tool in FSL^68,69^. The intensity-based mask selected voxels with signal intensity greater than four times the mean brain signal, leveraging the bright CSF contrast in T_2_-weighted magnitude images. Within the CSF mask, spatial smoothing was applied using a Gaussian kernel with a 3-mm full width at half maximum (FWHM). To avoid potential bias from phase wrapping induced by fast flow, voxels exhibiting large phase corresponding to velocities exceeding ±1280 μm/s (80% of the VENC) at any point of the time series were excluded from both analysis and visualization. Retrospective cardiac gating and net flow calculations were then performed using the full time series. For cardiac gating, velocity maps were binned into cardiac phases based on recorded physiological signals and acquisition timing, and averaged within each bin. For net flow estimation, two approaches, time averaging of velocities without cardiac-gating and across the cardiac-gated time series, were used and compared to obtain the temporally averaged mean velocity for each voxel. Three-directional velocity components (x, y, z) were combined to generate 3D flow-vector-fields representing both flow direction and magnitude.

For group-level processing (**Fig. 1c**), CSF flow velocity maps from individual subjects were registered to their corresponding MPRAGE image using FreeSurfer^70^ and then spatially normalized to MNI space using the 1-mm T^1^-weighted template and the Advanced Normalization Tools (ANTs) software^71,72^. To preserve flow directionality after spatial normalization, voxel-wise flow vector orientations were corrected using the rotational component of the Jacobian matrix derived from the deformation fields. Finally, velocity maps were averaged across subjects for each encoding direction, and the three directional components were combined to generate group-averaged CSF flow-vector-field maps for both cardiac-gated and net flow analyses. Combined velocity magnitude across all three directions were computed as the root-sum-of-squares (RSS) of the velocities from the three encoding directions at each voxel. Velocity range was calculated as the difference between maximum and minimum velocities across the cardiac cycle for each encoding direction, and the three directional ranges were combined using RSS for each voxel. Peak velocity was defined as the maximum absolute velocity observed across the cardiac cycle, and the peak cardiac phase was determined as the cardiac phase corresponding to the peak velocity.

### Data visualization and analysis

Velocity maps (e.g., combined velocity, velocity range, peak velocity) and other data such as peak-phase maps and vessel probability atlas overlaid on the anatomical T^1^-weighted MPRAGE image were visualized in Freeview^70,73,74^. Three-dimensional CSF flow-vector-fields were visualized in ParaView (Kitware, Inc.) using the “Glyph” representation with “arrow” geometry to represent the voxel-wise vector. The vector orientation indicated flow direction, vector length was proportional to 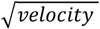, and color encoding represented A-P and S-I flow directions. A velocity threshold of 10 μm/s was applied to exclude low-magnitude vectors in the flow-vector-field maps. Flow-direction polarity was defined consistently across analyses, with the x-axis corresponding to left–right (positive left-to-right), the y-axis to posterior–anterior (positive posterior-to-anterior), and the z-axis to inferior–superior (positive inferior-to-superior).

For segmentation of periarterial and non-periarterial subarachnoid CSF, regions within five voxels of major arteries—defined using the vessel probability atlas with values greater than 5%—were classified as periarterial space, while the remaining subarachnoid regions were categorized as non-periarterial. For surface-based CSF ROI analysis, the T1-w MPRAGE data of each subject were segmented using FreeSurfer^70,73^ to generate cortical and subarachnoid space ROIs. The ‘DKTatlas’ was used for cortical parcellation, and the resulting cortical parcels were extended outward from the cortical gray matter into the adjacent CSF space to define subarachnoid space ROIs. This yielded 70 cortical-based subarachnoid space ROIs for each subject, which were used to compute ROI-based flow metrics, including velocity range over a cardiac cycle, net flow velocity, and 3-directional cardiac-gated and net flow, for repeatability analysis. Within each ROI, the mean parameter value was calculated from the top 6000 voxels with the highest magnitude. For statistical analysis, scan–rescan repeatability was evaluated using Pearson’s correlation coefficients calculated by pooling all participants and ROIs using MATLAB (MathWorks) ‘corrcoef’ function, and intraclass correlation coefficients (ICC; two-way mixed-effects model, absolute agreement) across all participants computed for each ROI. Intra- and inter-subject variability were quantified using the coefficient of variation (CV). Subject-averaged ROI-based flow metrics were overlaid onto the reconstructed cortical surface for visualization.

## ACKNOWLEDGMENTS

This work was supported by the National Institutes of Health (grants R00-AG083056, U24-NS129893, R01-EB036507, R01-AT011429, U19-NS128613, R01-AG070135, P41-EB030006, S10-OD023637).

## Author contributions

Conceptualization: ZD, FW; Methodology: ZD, FW; Investigation: ZD, FW, LDL, TGR; Supervision: ZD, FW, LDL, LLW, BRR; Writing—original draft: ZD, FW; Writing—review & editing: ZD, FW, LDL, TGR, LLW, BRR

## Extended Data

**Extended Data Fig. 1.**
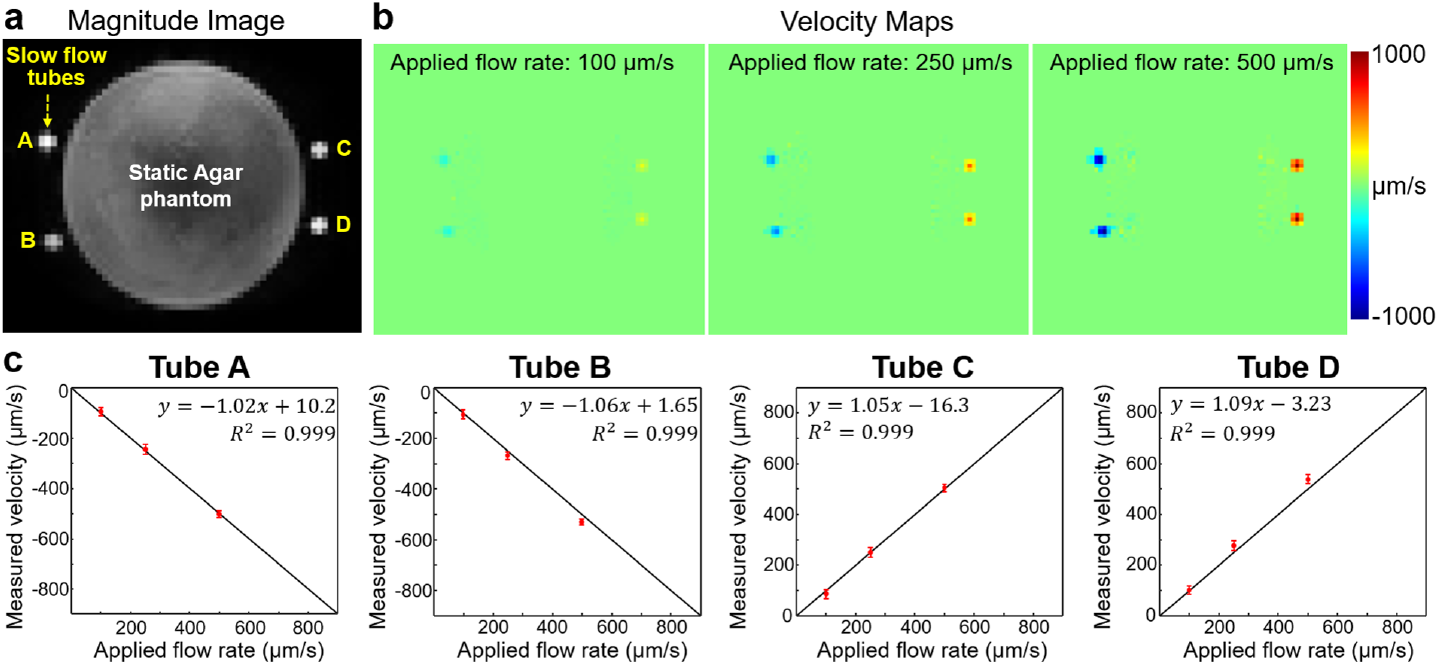
Slow flow phantom validation. **a**, Magnitude image of the phantom, showing four tubes (A-D) containing slow flow positioned around a static agar phantom. Flow was oriented along the z direction, with opposite flow directions in tubes A and B compared with tubes C and D. **b**, Measured velocity maps showing values consistent with the three reference applied flow rates of 100, 250, and 500 μm/s. **c**, Scatter plots (mean ± standard deviation) of measured versus applied velocities in the four tubes. In each plot, each data point is shown as a dot with an error bar representing the mean and standard deviation across time frames, and an identity line is shown in black. Fitted parameters are shown at the top of each plot. Measured velocities exhibited strong agreement with the applied velocities, with Pearson’s correlation coefficients > 0.99 for all tubes.

**Extended Data Fig. 2.**
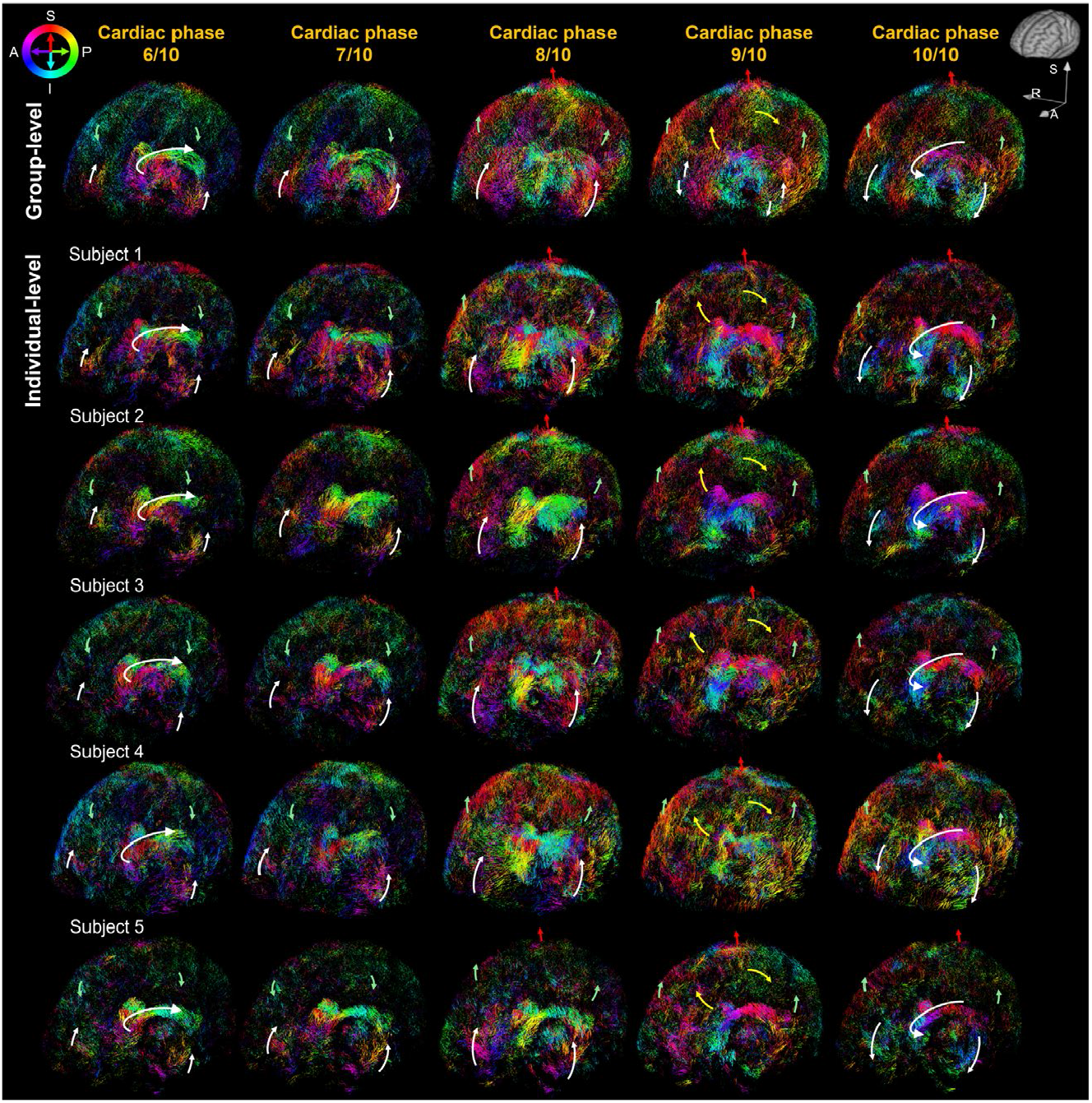
Individual-level brain-wide spatiotemporal subarachnoid CSF flow patterns (cardiac-gated). Individual cardiac-gated CSF flow-vector-field maps at representative cardiac phases, shown as whole-CSF-volume projections from angled sagittal views, illustrating large-scale and localized flow patterns highlighted by colored arrows. White arrows: lateral ventricle and Sylvian fissure; yellow arrows: interhemispheric fissure; green arrows: outer convexity subarachnoid space; red arrow: subarachnoid space near the SSS. Group-level cardiac-gated CSF flow-vector-field maps are also shown on the top row.

**Extended Data Fig. 3.**
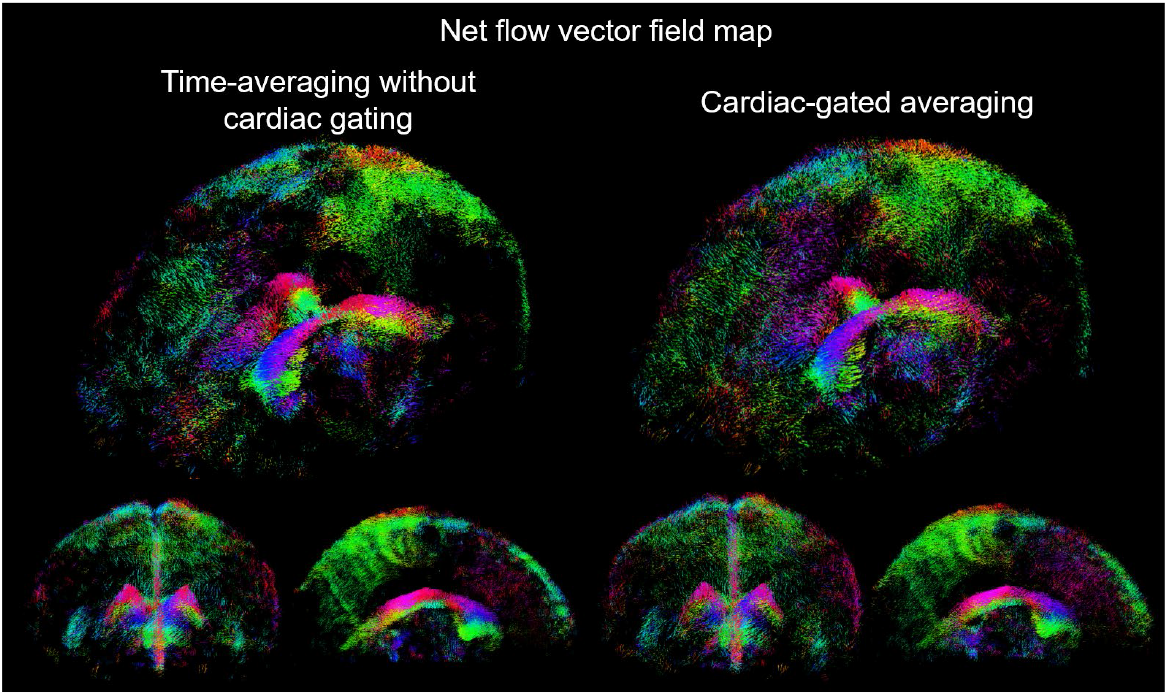
Net flow-vector-field maps computed using time-averaging without cardiac gating and cardiac-gated averaging. Whole-CSF-volume projections of the net flow-vector-fields from angled sagittal, coronal and sagittal views are shown. Net flow-vector-field maps derived from time-averaging without cardiac gating and from averaging cardiac-gated data (binned into 10 cardiac phases) exhibited similar flow patterns.

**Extended Data Fig. 4.**
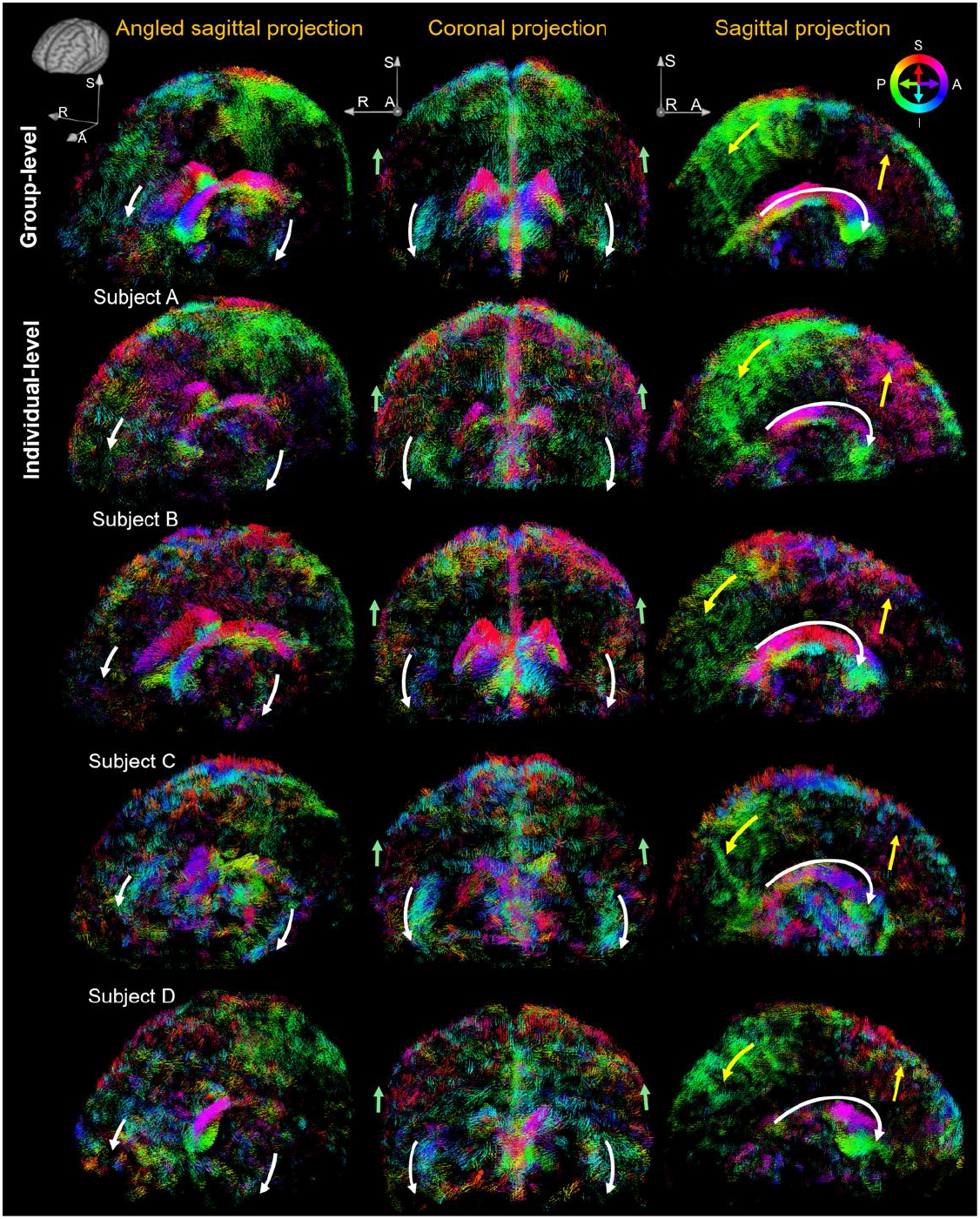
Individual-level brain-wide spatiotemporal subarachnoid CSF flow patterns (net). Individual net CSF flow-vector-field maps, shown as whole-CSF-volume projections from angled sagittal, coronal, and sagittal views, illustrating flow patterns highlighted by colored arrows. White arrows: lateral ventricle and Sylvian fissure; yellow arrows: interhemispheric fissure; green arrows: outer convexity subarachnoid space. Group-level net CSF flow-vector-field maps are also shown on the top row.

**Extended Data Fig. 5.**
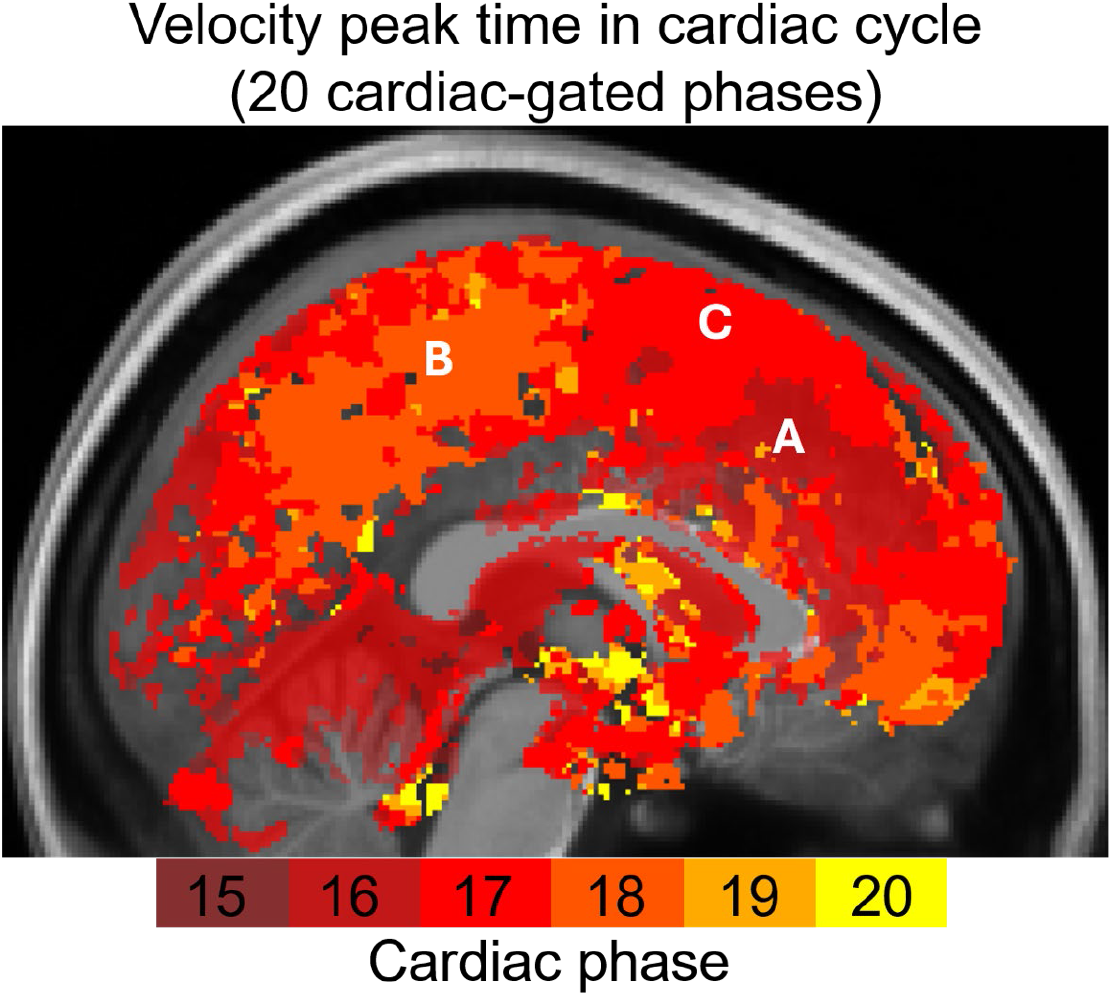
Velocity peak time map derived from 20 cardiac-gated phases. The finer cardiac-phase resolution reveals temporal differences within the interhemispheric fissure, with the anterior segment (A) reaching peak velocity at cardiac phase 16, followed by the middle segment (C) at phase 17 and the posterior segment (B) at phase 18, indicating spatiotemporal propagation of the pulsation-driven flow.

## Notes

### Competing Interest Statement

The authors have declared no competing interest.

