## Supplementary figures and images for "Mapping subarachnoid cerebrospinal fluid circulation in the human brain"

### Supplementary Video 1

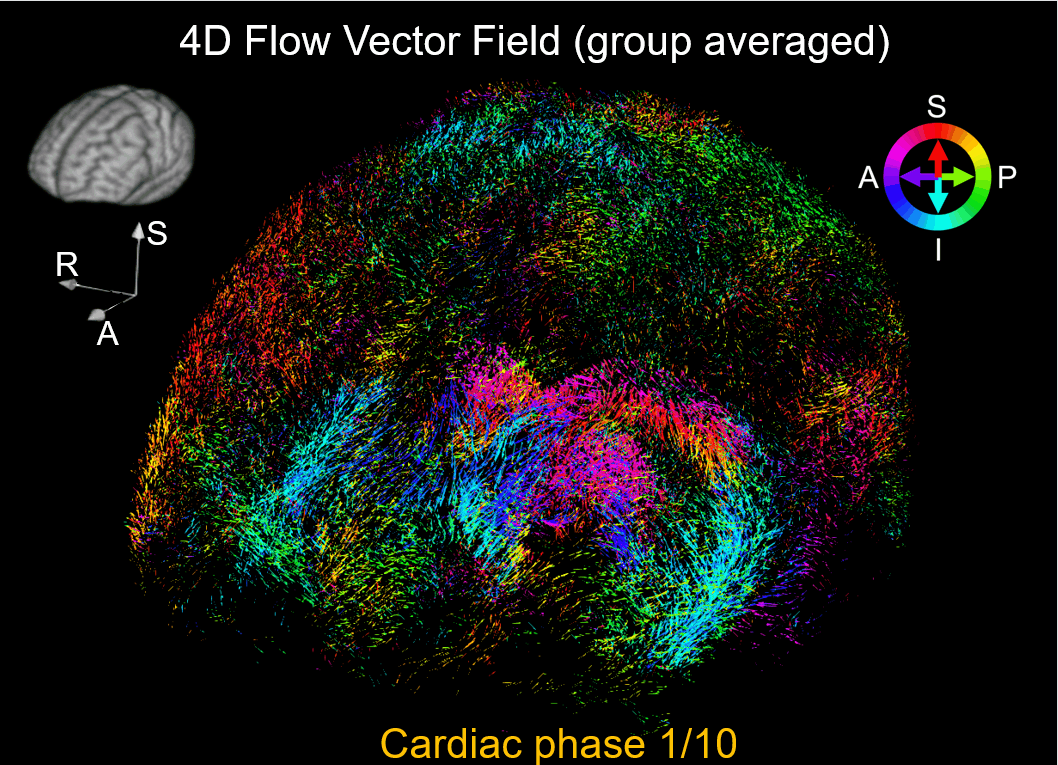
